# Validating Regulatory Predictions from Diverse Bacteria with Mutant Fitness Data

**DOI:** 10.1101/091405

**Authors:** Shiori Sagawa, Morgan N. Price, Adam M. Deutschbauer, Adam P. Arkin

## Abstract

Although transcriptional regulation is fundamental to understanding bacterial physiology, the targets of most bacterial transcription factors are not known. Comparative genomics has been used to identify likely targets of some of these transcription factors, but these predictions typically lack experimental support. Here, we used mutant fitness data, which measures the importance of each gene for a bacterium’s growth across many conditions, to validate regulatory predictions from RegPrecise, a curated collection of comparative genomics predictions. Because characterized transcription factors often have correlated fitness with one of their targets (either positively or negatively), correlated fitness patterns provide support for the comparative genomics predictions. At a false discovery rate of 3%, we identified significant cofitness for at least one target of 158 TFs in 107 ortholog groups and from 24 bacteria. Thus, high-throughput genetics can be used to identify a high-confidence subset of the sequence-based regulatory predictions.

## Introduction

Gene regulation is a key mechanism used by bacteria to survive in varying environments. The major class of regulators in prokaryotes are transcription factors (TFs), which are proteins that increase or decrease the transcription of their target genes. Transcription factors target specific genes by binding to specific sites near those genes’ promoters. Each transcription factor has a specific pattern of DNA sequences that it binds to, called a motif (Rodionov, 2007).

Since the typical bacterial genome encodes over 100 TFs whose targets and functions are not known (Charoensawan et al., 2010), it is critical to study TFs at large scales. Experimentally, TF-target interactions have been predicted by using high-throughput co-expression data (Faith et al., 2007) or by using chromatin immunoprecipitation and related techniques to identify a TF’s binding locations on the genome (Ishihama et al., 2016). However, co-expression is indirect evidence of a regulatory relationship (Rodionov, 2007) and these predictions are incomplete and have high false positive rates (Faith et al., 2007). For instance, the context likelihood of relatedness algorithm identified 11% of known interactions at a false positive rate of 40% in *E. coli* (Faith et al., 2007). Location data also has several limitations. First, location data fails to detect some interactions due to inefficient cross-linking at some locations (Rodionov, 2007) or poor TF binding in tested conditions. Second, location data may not imply regulatory relationships, as it may detect binding that does not affect expression (Wu et al., 2007). Lastly, while genome-wide coexpression is available for over 20 species (Meysman et al., 2013; Faith et al., 2007; Wendisch, 2003), to our knowledge large-scale location data is only available for a few bacteria (Ishihama et al., 2016; Galagan et al., 2012; Rajeev et al., 2011).

Alternatively, TF-target pairs have been predicted using comparative genomics. Comparative genomics methods rely on a key assumption that transcription factor binding sites are more conserved than other intergenic sequences. For example, a known motif for a TF can be used to identify targets of the TF’s orthologs by searching for occurrences of the known motif (Rodionov, 2007), using tools such as Patser (Stormo et al., 1989). The identified genes are predicted to be regulated by an ortholog of the TF, especially if multiple orthologous targets have the site. Another approach is to predict new motifs and to search for their associated targets. In order to predict a motif, a training set, composed of upstream regions of potentially co-regulated genes, is collected from multiple orthologous genomes (Rodionov, 2007). Then, a conserved sequence is identified in the training set using motif discovery algorithms such as MEME (Bailey and Elkan, 1994). The motif is validated if it is conserved across orthologous genomes. A major challenge in this approach is the association of a motif to the correct TF (Rodionov, 2007). There are many heuristics for predicting the TF-motif pairs (Tan et al., 2005), but experimental evidence can significantly improve the accuracy (Rodionov, 2007).

RegPrecise provides a manually-curated collection of comparative genomics predictions that associate TFs with motifs and targets (Novichkov et al., 2013). These predictions are based on both known and newly predicted motifs. Training sets for motif predictions are mainly composed of genes in a functional pathway (Novichkov et al., 2010). Most RegPrecise predictions lack experimental evidence.

Here, we consider the utility of mutant fitness data for validating the computational predictions from RegPrecise. To collect the fitness data, transposon mutant strains, each of which has one transposon insertion in the genome, are grown in a pool of ~40,000-500,000 strains in various conditions. The effect of mutating each gene on growth in each condition is then measured by quantifying the change in abundance of their mutant strains (Wetmore et al., 2015). We used fitness data for most genes in 25 bacteria across many conditions (Price et al., 2016; http://fit.genomics.lbl.gov/). The fitness patterns of activators and repressors are expected to be positively and negatively correlated with the fitness patterns of their targets (Deutschbauer et al., 2011).

In this study, we show that many characterized transcription factors have strongly correlated fitness patterns with a small subset of their targets. As a consequence, we consider a TF-motif association from comparative genomics to be validated if at least one target’s fitness pattern is significantly correlated with the TF’s pattern. In RegPrecise, we validated 158 TFs in 107 ortholog groups with a 3.0% false discovery rate (FDR). We also classified most of these TFs as activators (124) or repressors (28) based on the sign of their cofitness with their target genes. Thus, the mutant fitness data provides experimental support for the roles of over 100 TFs that were previously predicted by comparative genomics.

## Results

### Cofitness of transcription factors and their targets

Activating TF-target pairs are expected to have strong positive cofitness, where cofitness is the correlation between the fitness patterns of two genes. If a transposon is inserted into an activator, then the target gene’s expression would be reduced. If a transposon is inserted into a target gene, then there would no longer be functional transcripts of the target gene. Since the amount of functional transcript of the target is reduced under both scenarios, the activator and its targets are expected to have positively correlated fitness patterns. In contrast, repressors are expected to be anti-cofit (have negative cofitness) with their targets. If a transposon is inserted into a repressor, then the target gene expression would be elevated. If a transposon is inserted into a target gene, then there would no longer be functional transcripts of the target gene. If elevated and reduced levels of functional target transcripts lead to opposite phenotypes, then the two fitness patterns are expected to be negatively correlated. Similarly, TF-target pairs are expected to have extreme ranks in cofitness and anti-cofitness. Cofitness rank is the rank of TF-target cofitness relative to the TF’s cofitness with other genes in the genome, where the most positive cofitness has the top rank. Anti-cofitness rank is the ranking in reverse order.

In *E. coli*, transcription factors do tend to be highly cofit or anti-cofit with at least one of their experimentally-identified targets. For example, DcuR is an activating transcription factor that regulates genes involved in C4-dicarboxylate metabolism, and it has high cofitness with one of its targets, *dctA*. As shown in Figure 1A, *dcuR* and *dctA* are important for aerobic growth with dicarboxylates (malate or succinate) as the carbon source, as expected. More importantly, *dcuR*’s fitness pattern closely matches the fitness pattern of *dctA* (Figure 1A), with cofitness of 0.97 and rank 2 out of 3,788 (Figure 1B,C). Other targets of DcuR have weak cofitness and ranks (Figure 1C). For example, fumarate reductase (*fdrABCD*) is only expected to be on important under anaerobic conditions, and its expression can be induced by another regulator besides DcuR (namely Fnr), so it’s not surprising that it has a different fitness pattern than *dcuR*. In contrast, *treR*, which represses genes involved in trehalose transport and degradation, has a fitness pattern that seems to be the opposite of the fitness pattern of its target, *treB.* As shown in Figure 1D, *treR* and its targets have strong phenotypes when trehalose is the carbon source, as expected. More importantly, *treR* and *treB* have strong negative cofitness of -0.59 and a top anti-cofitness rank of 1 (Figure 1C). TreR’s other target, *treC*, has weak cofitness and rank (Figure 1C,D); even though *treBC* forms an operon, the fitness pattern of *treC* seems to be more complex because, aside from its role in trehalose utilization, TreC can modify maltose (Decker et al., 1999).

**Figure 1:**
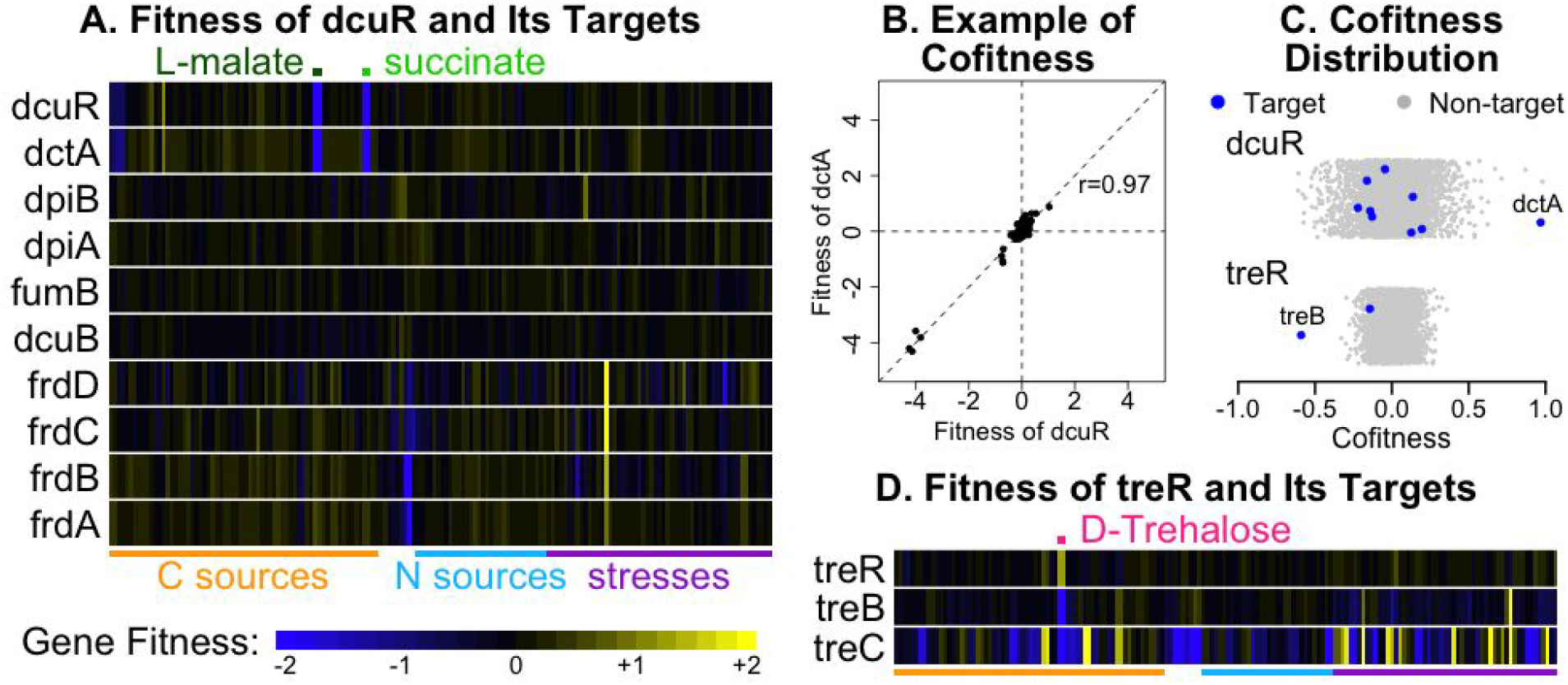
Examples of fitness patterns of experimentally-identified TF-target pairs in *E. coli*. (A) Heatmap showing the fitness values of the activator *dcuR* and its targets across 162 experiments. Gene fitness is a log_2_ ratio that describes the change in abundance of strains with a transposon inside the gene over the course of pooled incubation (Wetmore et al., 2015). Positive fitness suggests that the gene is detrimental to growth in the condition, while negative fitness suggests that the gene is important for growth. The labels above and below the heatmap describe the growth conditions. (B) Scatter plot of the fitness values of *dcuR* and *dctA*. Lines show *x=0*, *y=0*, and *x=y*. (C) Distribution of cofitness values for *dcuR* and *treR*. Each panel shows the cofitness of the transcription factor with 3,788 genes in the *E. coli* genome that have fitness data (x axis). The *y* axis is random. (D) Heatmap showing the fitness values of the *treR* repressor and its targets across 162 experiments.

More broadly, we examined the cofitness of all TF-target pairs that have been identified experimentally in E. coli (Salgado, 2013). TFs that are both activators and repressors were excluded to avoid ambiguous cases, which left 44 activating TFs and 57 repressing TFs. As shown in Figure 2A, TF-target pairs are highly enriched in strong positive cofitness (or anti-cofitness) relative to shuffled pairs. For example, cofitness above 0.5 is enriched in activating TF-target pairs by 15-fold, while anti-cofitness under -0.5 is enriched in repressing TF-target pairs by over 21-fold. Figure 2B shows that cofitness ranks of 1-50 are enriched in activating pairs by 10-fold, while anti-cofitness ranks of 1-50 are enriched in repressors by 4-fold.

**Figure 2:**
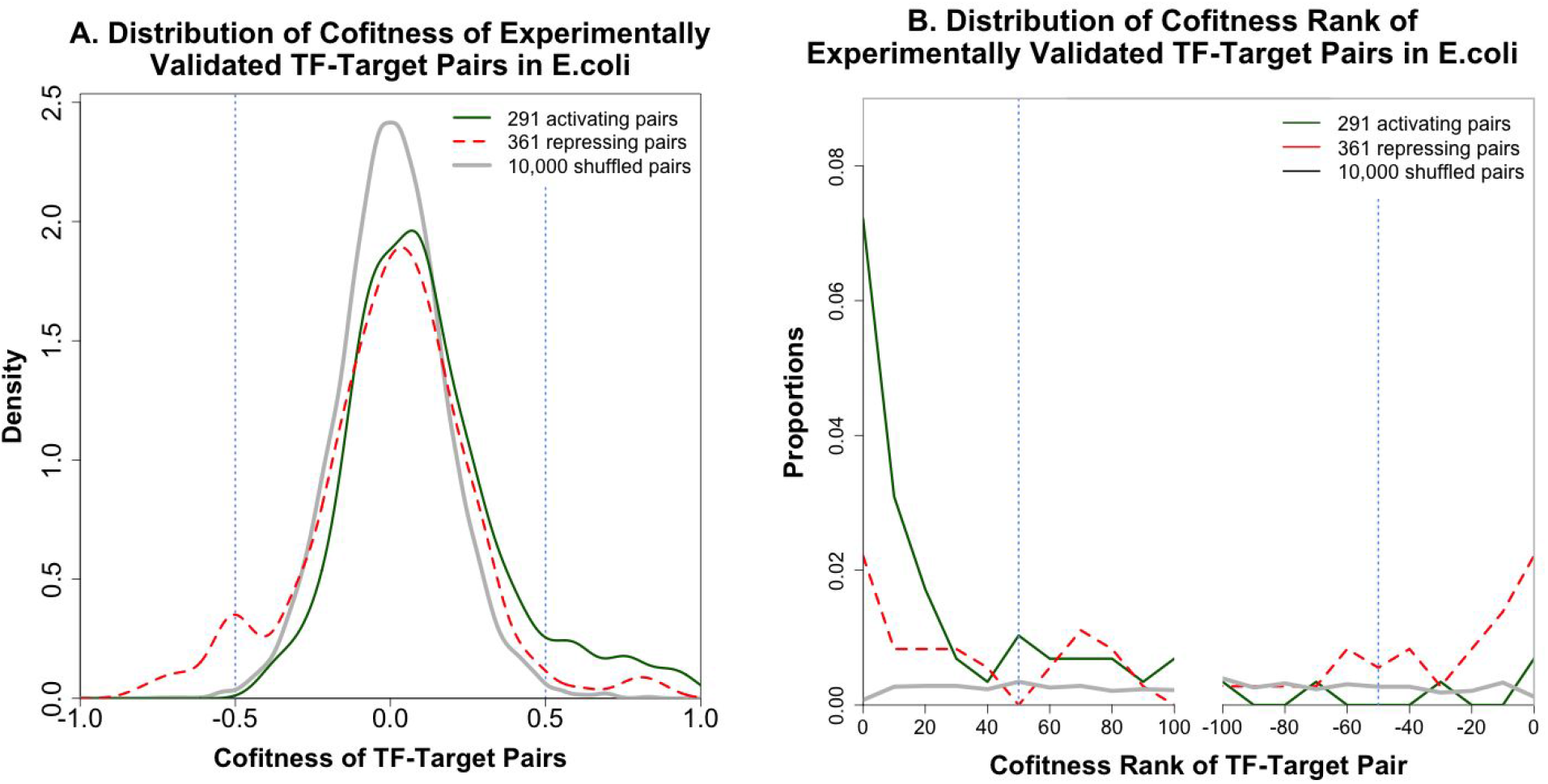
Distribution of cofitness and cofitness ranks in experimentally-identified TF-target pairs in *E. coli*. (A) The distribution of cofitness values for activating TF-target pairs, repressing TF-target pairs, and shuffled pairs. The distributions were smoothed using gaussian kernel density estimates. (B) The distribution of cofitness ranks for activating TF-target pairs, repressing TF-target pairs, and shuffled pairs. On the right side, ranks for anti-cofitness are shown with negative numbers, so that the most anti-cofit pair has *x = -1*. The proportion of pairs within each window of 10 cofitness ranks is plotted.

### Statistical Approach

Applying the above observations, we formulated a method to validate TF-motif predictions from comparative genomics. It is useful to test these predictions at the motif level for two reasons. First, the predictions are motif-based, and the association of a TF with a motif is a major source of uncertainty in comparative genomics predictions. Second, the transcription factor may be cofit with a subset of its targets, as we observed for the characterized regulators *dcuR* and *treR*. Figure 3 demonstrates this trend for several other characterized transcription factors from *E. coli*. For this reason, the method validates TF-motif predictions based on the predicted target that is most strongly cofit or anti-cofit with the TF.

The method uses cofitness rank in addition to cofitness, since the distribution of cofitness varies widely among TFs (Figure 3). In fact, the standard deviation of the cofitness distribution varies from 0.08 to 0.31 among the experimentally-characterized TFs in *E. coli*. As a result, it is important to look at the cofitness of the TF-target pairs relative to the TF’s cofitness with other genes. For example, in Figure 3, the cofitness value of 0.61 for *deoR* and its most cofit target seems significant, but such a cofitness would be less convincing for *metR*. Of the 97 characterized TFs in *E. coli* that have significant phenotypes (at a false discovery rate under 5%; Price et al., 2016), 64 (66%) have highly-ranked cofitness or anti-cofitness (rank 1-50) with at least one target.

**Figure 3:**
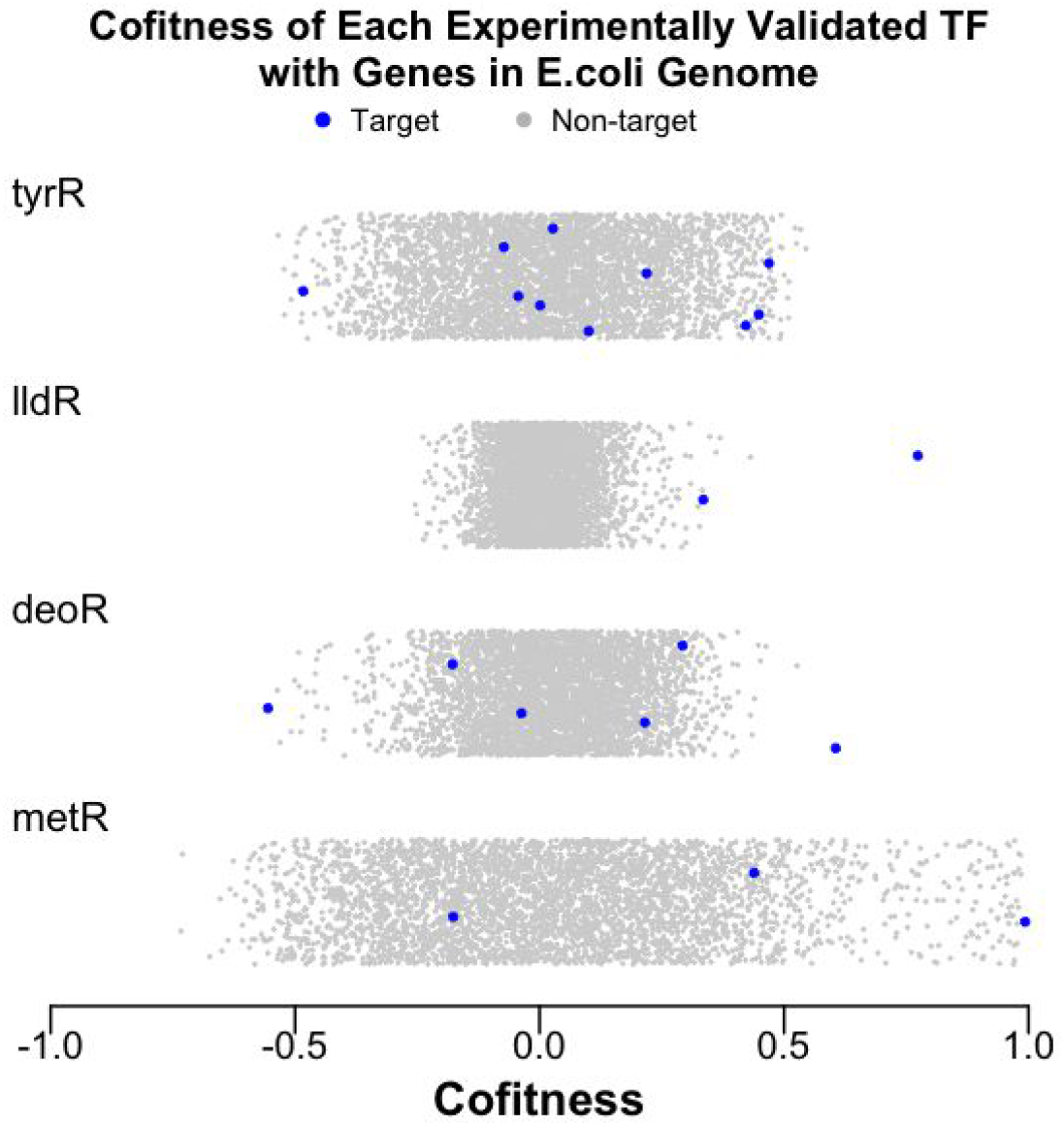
Distribution of cofitness values for four experimentally characterized transcription factors in *E. coli*. Each panel shows the cofitness of the transcription factor with all other 3,788 genes in the *E. coli* genome with fitness data (x axis). The y axis is random.

To validate each TF-motif association, we conducted a hypothesis test. The null and alternate hypotheses state that the the TF is not and is associated with the motif, respectively. Under the null hypothesis, any cofitness between the TF and a target is due to random chance; more precisely, the cofitness of the TF with its predicted targets is assumed to follow the same distribution as for a random sample of genes. We calculate two p-values. First, we calculate the probability that a random sample of *T* genes would have a top (anti-)cofitness rank as extreme as the observed value, where *T* is the number of TF’s targets not including the TF. Here, we use cofitness ranks that are corrected for a bias, in which closely located gene pairs tend to have slightly higher cofitness (see Methods). Secondly, to ensure that the value of cofitness is statistically significant, the Fisher transformation p-values for the most positive and negative cofitness values were computed. Only TFs that have a significant phenotype in at least one condition (false discovery rate under 5%; Price et al., 2016) were tested, and they were considered validated if both p-values were 0.01 or below.

### Validation of RegPrecise Predictions

The hypothesis test was applied to the RegPrecise predictions, but since predictions are not available for 17 of the 25 bacteria with fitness data, we tested orthologous predictions for those organisms. An orthologous prediction is a pair of genes that are orthologous to a TF and a target in an original prediction, where the motif of the original prediction exists near the target’s ortholog (see Methods). In RegPrecise, we tested 479 TFs and we validated at least one target for 158 TFs in 107 ortholog groups at a false discovery rate of 3.0%. (The FDR was estimated from the expected false positive rate * cases tested / positive cases = 0.01 * 479 / 158 = 0.030.) The validated predictions are listed in tables S1 and S2. Validated TFs are involved in a variety of pathways, as summarized in Figure 4.

**Figure 4:**
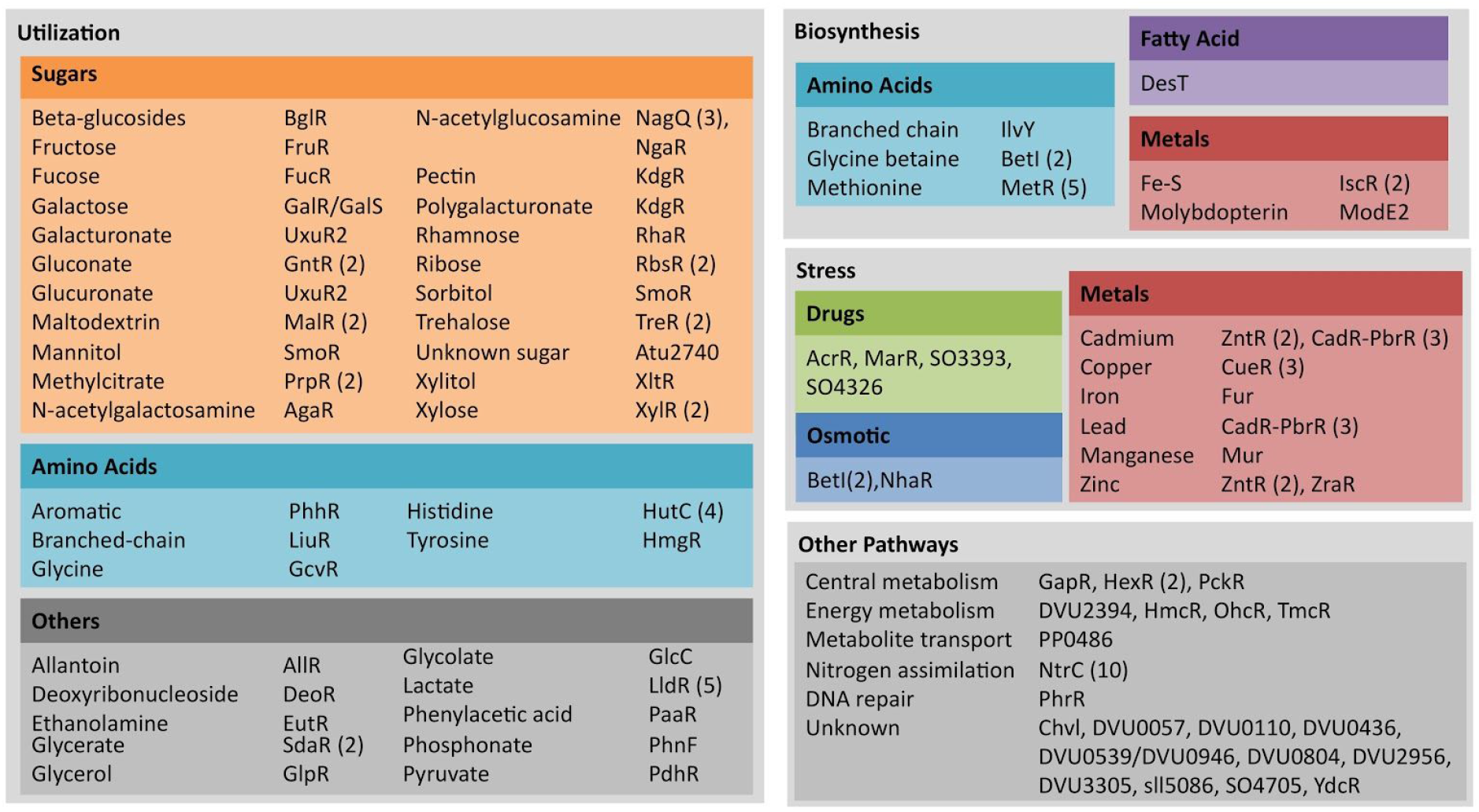
Summary of validated RegPrecise predictions. If multiple ortholog groups with the same TF name were validated, the number of validated ortholog groups is noted. An ortholog group is a group of orthologous TFs from closely related organisms with a conserved motif.

### Sources of False Positives

Although our statistical test is stringent enough to obtain a low rate of false positives due to random correlations, false positives could also arise from other sources. The most prominent source is polar effects, where an insertion in a gene leads to transcription termination and reduced expression of downstream genes. Polar effects on a gene would cause positive cofitness between the gene and its co-transcribed genes, leading to false validations by positive cofitness. In particular, if a TF is co-transcribed with another gene and does not regulate the operon, then the TF may be falsely validated due to polar effects. TF predictions that are potentially affected by polar effects are more likely to be validated due to positive cofitness than other predictions (36% vs 20%) and this difference is statistically significant (16%± 8%, 95% CI). As a control, we also examined the rate of validation by anti-cofitness and found no effect of co-transcription (a difference of -3%± 4%, 95% CI). Thus, a significant fraction of predictions that are potentially affected by polar effects may be falsely validated. However, analysis of known TFs in *E. coli* suggests that only a small subset of TFs experience significant polar effects. Of 14 pairs of *E. coli* TFs that have significant phenotypes and have a potential polar effect (have a co-transcribed adjacent gene that is not a target), five pairs had a cofitness rank of 1-50 (*ompR-envZ, yehT-yehU, dcuR-dcuS, basR-basS,* and *uhpA-uhpB*). However, since all five of these pairs are histidine kinase-response regulator pairs and are thus expected to be cofit, they do not necessarily indicate polar effects. Overall, for 65 of the positively validated TFs (50%), they or their orthologous TFs are validated by targets that are not in the same operon as the TF.

Secondly, complex regulatory networks could lead to false positives. For example, if a TF regulates another TF, the first TF may be cofit with the second TF. This cofitness might appear to validate the second TF with the first TF as a target. Alternatively, the first TF might be cofit with targets of the second TF. We believe that such false positives should not be common since the majority of bacterial TFs do not regulate other TFs: 43% of characterized TFs in *E. coli* regulate another TF (Salgado et al., 2013). In addition, of 21 characterized TFs in *E. coli* that are “validated” by the fitness data and our statistical approach, only one TF is more cofit or anti-cofit with another TF than with their best target (*marR* is more cofit with *acrR*, which regulates *marR*).

### Prediction of Regulatory Sign

The method can also be used to predict the sign of the regulation. If validated due to cofitness, then the TF is most likely an activator. If validated due to anti-cofitness, then it is most likely a repressor. For known transcription factors in *E. coli*, this approach is mostly accurate. For this analysis, we excluded TFs that are both activators and repressors, as either prediction would be accurate. Of the 9 remaining TFs from *E. coli* that we “validated” by positive cofitness, 8 are activators. And all 3 of the remaining *E. coli* TFs that were “validated” by anti-cofitness are repressors. In addition, the misclassified regulator, PuuR, can be explained by polar effects. Lastly, there were 5 ambiguous cases (AcrR, CytR, DeoR, DsdC, GlcC), where the TF was “validated” for both cofitness and anti-cofitness. This seems to happen because two target genes have negatively-correlated phenotypes, so that the TF is cofit with one target and anti-cofit with the other.

Using this approach, we predicted the signs of validated TFs from RegPrecise: 124 activators, 28 repressors, and 6 ambiguous. In some cases, the sign suggested by the cofitness data conflicts the annotation in RegPrecise. For example, MalR, which is in the LacI family, is annotated as a repressor. To our knowledge, there is no experimental evidence of the regulational sign of MalR or its orthologs. However, *malR*’s fitness pattern is strongly correlated with the fitness pattern of *malT* (*Shew_1874*) in *Shewanella loihica* with cofitness of 0.94 and rank of 1, as demonstrated in Figure 5. Similar fitness patterns are also seen in *Shewanella* sp. ANA-3 and *Shewanella amazonensis*. Predictions in all three organisms are not affected by polar effects. In addition, MalR binds over 300 basepairs upstream of the transcription start site of its target, *malT* (Shao et al., 2014), as is often observed with activators (Rodionov, 2007). Thus, it seems likely that the regulational sign of MalR is misannotated in RegPrecise, and it is actually an activator, at least in the presence of maltose. Fitness data also suggest that SO4705 (from *Shewanella oneidensis*, *Shewanella* sp. ANA-3, *Shewanella amazonensis*), PckR (from *Sinorhizobium meliloti*), SdaR (from *Shewanella oneidensis*), and ModE2 (from *Desulfovibrio vulgaris*) are activators, contrary to their annotations. These TFs are not affected by polar effects. However, for SO4705, the binding sites are very close to or downstream of the transcription start sites of the targets (Shao et al., 2014), which suggests that SO4705 may yet be a repressor.

**Figure 5:**
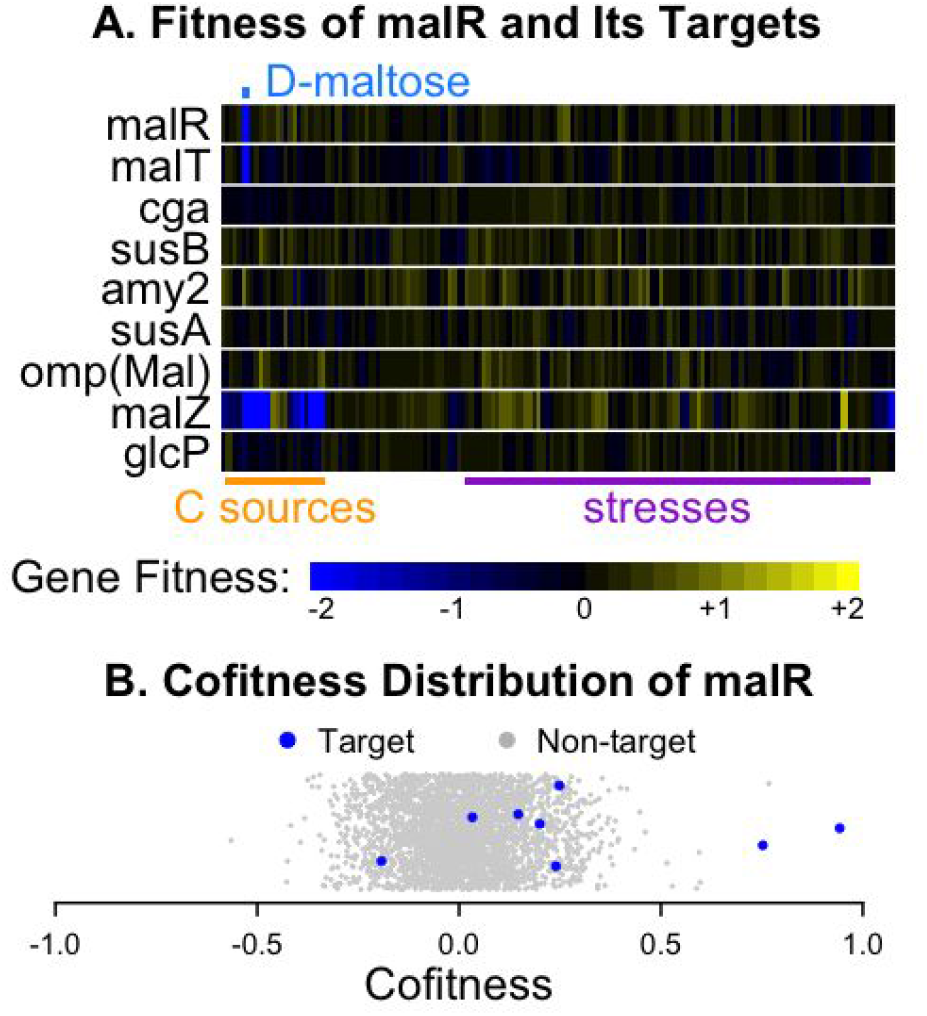
Fitness pattern of *malR* in *Shewanella loihica*. (A) Heatmap showing fitness values of *malR* and its targets across 160 experiments. The labels above and below the heatmap are conditions. (B) Distribution of *malR*’s cofitness with all other 3,008 genes in the *Shewanella loihica* genome with fitness data (x axis). The y axis is random.

### Application of the Approach to Coexpression Data

To our knowledge, coexpression has not been used to validate regulatory predictions in bacteria at a large scale. We tested our approach on coexpression data from *E. coli* and *Shewanella oneidensis* in the Many Microbe Microarrays Database (Faith et al., 2008). First, we tested characterized TFs in *E. coli* with at least one target that’s not co-transcribed with the TF. We “validated” these TFs based on their coexpression with a target in a different operon. Of 171 such TFs with coexpression data, 16 of them were validated by coexpression with a false discovery rate (FDR) of 10.6%. For comparison, of 89 TFs with a significant phenotype, 17 were validated by cofitness at a 5.2% FDR. Of the 89 TFs with both a significant phenotype and coexpression data, only four (*dsdC, galS, malT,* and *puuR*) are validated by both methods, which suggests that the two validation methods are complementary. Second, we tested our approach on TFs from *Shewanella oneidensis* that have predictions from RegPrecise. Of 50 TFs tested, we validated 5 TFs at 10.0% FDR.

Coexpression seems to give a weaker signal than cofitness, as indicated by the higher FDRs. TFs often lack strong coexpression with any of their targets. For example, of 13 characterized TFs in *E. coli* that were “validated” by cofitness but not by coexpression, 12 TFs have coexpression of magnitude under 0.5 with all targets. *fucR* has moderate coexpression (0.52) and coexpression rank (37) with its target *fucA*, but these were not strong enough for validation. Overall, although validation by coexpression has a higher FDR, coexpression can be used to validate predictions that are not validated by cofitness.

## Discussion

### Advantages and Limitations of Use of Fitness Data

Although fitness data successfully validates a subset of the regulatory predictions, it has several limitations compared to coexpression analysis. First, fitness data shows a correlation for a smaller fraction of TF-target pairs than expression data does (Deutschbauer et al., 2011). In particular, cofitness analysis seems to be less effective for repressors. Although the characterized regulatory relationships in *E. coli* are roughly equally split between activators and repressors, most of the regulators that we validated had positive cofitness with their validated targets (78%). It appears that activators are much more likely to be cofit with a target than repressors are.

Second, fitness data measures changes in gene expression indirectly through its effects on growth, while the expression data does so directly. Nevertheless, cofitness seems to provide a stronger signal than coexpression, albeit for only a subset of the TF-target pairs. This was illustrated by the lower false discovery rate for “validating” known regulatory relationships in *E. coli* by cofitness as compared to coexpression.

Third, it is not possible to validate each TF-target pair by cofitness analysis because a TF’s fitness pattern may be determined only by a small subset of targets. This makes it difficult to make novel TF-motif or TF-target predictions based solely on fitness data.

Fourth, given our statistical approach, we cannot validate TFs with over a certain number of targets. For example, for TFs with over 18 targets in *E. coli* (which has 3,789 genes with fitness data), even a (anti-)cofitness rank of 1 would yield p > 0.01. For this reason, the cofitness analysis provides more powerful evidence for local TFs, which represent the vast majority of TFs in bacteria (Martinez-Antonio and Collado-Vides, 2003; Moreno-Campuzano et al., 2006).

Despite these limitations, there are some advantages to using the fitness data. First, cofitness analysis should be more robust than coexpression analysis if a TF and a non-target gene are coregulated by other factors. In this case, the two genes will be coexpressed, but they probably won’t be cofit. Because the fitness data measures the effect of disabling each gene, regulation of the gene does not determine its fitness pattern. Second, cofitness analysis is advantageous for validating co-transcribed TF-target pairs. If a TF and its potential targets are co-transcribed and coexpressed, it is unclear whether the coexpression is caused by co-transcription or by a regulatory relationship. But in the absence of a regulatory relationship, there should not be any cofitness (although because of polar effects, this will not always be true in practice). More broadly, the above two points illustrate that fitness data may have an advantage in identifying direct regulatory relationships because it relies on mutants of the TF. Finally, fitness data is complementary to other large-scale data. For example, coexpression profiles and cofitness data seem to have complementary signals, as they have been shown to be poorly correlated (Deutschbauer et al., 2011). Also, we found that in *E. coli*, there is little overlap between the characterized TFs that are “validated” by cofitness and by coexpression. The correlation may be poor because coexpression signals are expected to be weak for constitutively expressed TFs, while this is not expected for cofitness.

Fitness data may also be useful as a starting point for comparative genomics analyses. A significant fraction of putative TFs without comparative genomics predictions are cofit with at least one gene. For example, of 80 putative TFs (from the TF families in RegPrecise) that have no predictions in RegPrecise, 40 have significant phenotypes and of those, 24 have cofitness of over 0.8 or conserved cofitness of over 0.6, which are strong indications of a functional relationship (Price et al., 2016). Another 3 TFs had predictions that were not validated and have strong cofitness with other genes. The associations for these 27 TFs are listed in supplementary Table S3. Although these putative pairs are not expected to be high-confidence at this stage, further analysis using comparative genomics could lead to high-confidence predictions.

### Conclusions

Activating (or repressing) TFs tend to be cofit (or anti-cofit) with a subset of their targets. Based on this observation, we formulated a statistical method to validate TF-motif associations from comparative genomics predictions. The method was applied to RegPrecise predictions and 158 TFs in 107 ortholog groups were validated with a false discovery rate of 3.0%. Most of these TFs were predicted as activators (124) or repressors (28) based on the sign of the cofitness. Although the statistical test has a low rate of false positives due to random correlations, false positives could also arise from other sources, most notably polar effects. Nevertheless, validation by cofitness has several advantages over and is complementary to the use of other large-scale data.

## Methods

### Data

We obtained known TF-target pairs in *E. coli* from RegulonDB version 9.1, downloaded on April 7, 2015 (Salgado et al., 2013). Version 4.0 of RegPrecise was downloaded on April 29, 2016 (Novichkov et al., 2013). In RegPrecise predictions, each ortholog group has a unique regulog ID. Fitness data was obtained from http://genomics.lbl.gov/supplemental/bigfit/ and data from April 29, 2016 was used (Price et al., 2016). Coexpression data was obtained from the Many Microbe Microarrays Database on July 7, 2015 (Faith et al., 2008).

### Propagating Regulatory Predictions to Orthologs

Predictions were propagated from each organism with RegPrecise predictions to each organism with fitness data. In order to propagate predictions from one organism to another, we ran ublast (Edgar, 2010) between protein sequences of their genomes in both directions with thresholds *E* ≤ 10^-5^, identity ≥ 50%, and coverage ≥ 80%. A gene was considered a potential ortholog of another gene if they were best-scoring hits of each other. Orthologous TF-target pairs are pairs of genes where one gene is an ortholog of the TF and another is an ortholog of the target. If a TF of interest had multiple orthologs with predictions in RegPrecise, we chose just one bacterium to propagate from. We considered only bacteria that had at least one orthologous-TF pair (that is, there was at least one target with an ortholog in the bacterium of interest), and among those, we chose the bacterium whose TF was the most similar to the TF of interest (i.e., the highest bit score). Then, for each orthologous TF-target pair, we searched for the motif of the original TF around the target using Patser (Stormo et al., 1989). We identified the best-scoring hit found within 250 bp of the upstream end of each potential operon that the target ortholog belongs to, where potential operons were any series of adjacent genes that are on the same strand and less than 250 bp apart. A p-value for the motif hit was calculated by identifying the best motif hit for every gene in the genome and computing the fraction of genes that score at least as well as the target; genes in the same potential operon as the target ortholog were excluded. An orthologous TF-target pair was considered a propagated prediction if there is a motif hit with p≤0.01. There were significant motif hits for 56% of orthologous TF-target pairs from RegPrecise.

### Cofitness Bias Correction

Pairs of closely located genes tend to have slightly higher cofitness values than gene pairs that are far apart. This bias might arise from the effect of chromosomal position on DNA copy number; the fitness values are normalized to this trend (Wetmore et al., 2015), but a small bias may remain. Since TFs are often near their targets, cofitness values of TF-target pairs were corrected to account for this bias. To estimate the bias for each genome, we sampled 1,000 gene pairs that are at most 20KB apart and 1,000 gene pairs that are at least 50KB apart or are on a different scaffold. The median cofitness value for each set was computed. If the closely located pairs have higher median cofitness than the pairs that are far apart, the bias was estimated to be the difference in the median fitness. Otherwise, the bias was estimated as 0. The median and maximum bias among all 25 bacteria were 0.02 and 0.07. We corrected for the bias in positively cofit TF-target pairs that are at most 20 kb apart by subtracting the bias from the original cofitness value. The cofitness rank was calculated based on this corrected cofitness.

### Hypothesis Testing

The rank-based p-value quantifies the probability of getting a cofitness rank or an anti-cofitness rank as extreme as the observed value, given the number of targets. For each transcription factor, the rank-based p-values for activators and repressors were calculated as follows:

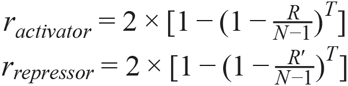

where N is the number of genes with fitness data in the genome, R is the top cofitness rank of the targets, R’ is the top anti-cofitness rank, and T is the number of targets not including the TF. The above p-values were calculated assuming sampling with replacement. However, their values are very close to the corresponding probabilities assuming sampling without replacement since T<<N (N is 1,899 to 6,384). Since we tested for two signs, rank-based p-values include the factor of 2 for Bonferroni correction. In addition, to ensure that the value of cofitness is statistically significant, the Fisher transformation p-values for the most positive and negative cofitness values were computed. These were corrected for testing across multiple target genes by Bonferroni correction. Finally, the overall p-values for activators and repressors are calculated as follows:

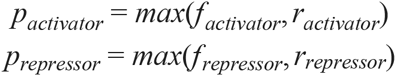

where *f_activator_* and *f_repressor_* are Fisher transformation p-values. We used a threshold of p≤0.01.

Only TFs with significant phenotypes (Price et al., 2016) were tested. In most bacteria, a gene has a significant phenotype if |fitness|>0.5 and |t|>4 for some condition. These thresholds are adjusted upwards as needed to ensure that the FDR for detecting significant phenotypes is under 5% in each bacterium (Price et al., 2016).

### Analysis of Polar Effects

A prediction whose TF is co-transcribed with a putative target is potentially affected by polar effects. Validation rates were compared between predictions with potential polar effects and others. Only TFs with a significant phenotype and at least one target with fitness data were considered. We computed the fraction of validated TFs in the two sets of predictions and used the two-proportion z-test to construct a 95% confidence interval for the difference.

### Putative TFs and Potential Targets

Putative TFs were identified from genes that belong to the same TIGRFAMs as TFs in RegPrecise predictions (Haft et al., 2012). (The list of putative TFs is not complete.) TIGRFAMs without DNA binding domains were manually filtered out in the analysis. Across the 25 genomes, we found 80 unpredicted putative TFs. We then sought to identify a potential target for each putative TF. We considered all genes that the putative TFs are not predicted to regulate in RegPrecise and the orthologous predictions. The potential target is the gene with the highest cofitness with the putative TF, out of genes that have conserved cofitness > 0.6 and conserved cofitness rank ≤ 10 with the putative TF, or that have cofitness > 0.8. (Conserved cofitness > 0.6 if the two genes have cofitness > 0.6 and an orthologous pair also has cofitness > 0.6.) Otherwise, a potential target was not identified for the putative TF.

### Statistical Tools

Statistical analyses were done with R version 3.1.1. Cofitness distributions were smoothed using the density function with default parameters. Code is available at http://genomics.lbl.gov/supplemental/cofittf/.

## Acknowledgements

This material by ENIGMA-Ecosystems and Networks Integrated with Genes and Molecular Assemblies (http://enigma.lbl.gov), a Scientific Focus Area Program at Lawrence Berkeley National Laboratory is based upon work supported by the U.S. Department of Energy, Office of Science, Office of Biological & Environmental Research under contract number DE-AC02-05CH11231.

## Supplementary Materials

These supplementary tables are available at http://genomics.lbl.gov/supplemental/cofittf/.

**Table S1:** List of transcription factors validated in original predictions.

**Table S2:** List of transcription factors validated in orthologous predictions

**Table S3:** Potential TF-target pairs identified by cofitness.

